# StrainR2 accurately deconvolutes strain-level abundances in synthetic microbial communities

**DOI:** 10.1101/2024.08.08.607172

**Authors:** Kerim Heber, Shuchang Tian, Daniela Betancurt-Anzola, Heejung Koo, Jordan E. Bisanz

**Affiliations:** Department of Biochemistry and Molecular Biology, Pennsylvania State University, University Park, PA 16802, USA; One Health Microbiome Center, Huck Life Sciences Institute, University Park, PA 16802, USA

**Keywords:** Strain-resolved metagenomics, strain quantification, k-mers, synthetic consortia

## Abstract

**Background:** Synthetic microbial communities offer an opportunity to conduct reductionist research in tractable model systems. However, deriving abundances of highly related strains within these communities is currently unreliable. 16S rRNA gene sequencing does not resolve abundance at the strain level, standard methods for analysis of shotgun metagenomic sequencing do not account for ambiguous mapping between closely related strains, and other methods such as quantitative PCR (qPCR) scale poorly and are resource prohibitive for complex communities. We present StrainR2, which utilizes shotgun metagenomic sequencing paired with a k-mer-based normalization strategy to provide high accuracy strain-level abundances for all members of a synthetic community, provided their genomes.

**Results:** Both *in silico,* and using sequencing data derived from gnotobiotic mice colonized with a synthetic fecal microbiota, StrainR2 resolves strain abundances with greater accuracy than other tools utilizing shotgun metagenomic sequencing reads and can resolve complex mixtures of highly related strains. Through experimental validation and benchmarking, we demonstrate that StrainR2’s accuracy is comparable to that of qPCR on a subset of strains resolved using absolute quantification. Further, it is capable of scaling to communities of hundreds of strains and efficiently utilizes memory being capable of running both on personal computers and high-performance computing nodes.

**Conclusions:** Using shotgun metagenomic sequencing reads is a viable method for determining accurate strain-level abundances in synthetic communities using StrainR2.

## BACKGROUND

Most metagenomic tools are unable to quantitatively resolve strain-level abundances; however, variation at the strain level is a crucial determinant of microbiome function and host-microbe interactions [1–3]. For example, some *Escherichia coli* strains are pathogens causing severe diarrhea, while others are described as probiotics used in treating diarrhea [4]. Strains of the same species (clonal populations within the species often represented by a cultured isolate) share a small proportion of their genome referred to as the core genome. However, many of the genes which drive important phenotypes are found in the variable, or accessory, portion of the genome [5]. Understanding the role that intra-species (infraspecific) variation plays in microbiome function, and the competitive interactions within these species are crucial for advancing our knowledge of how complex microbial communities assemble and function.

Synthetic communities offer a powerful tool that balance experimental reductionism with a biologically relevant scale across multiple systems [6–10]. These experiments are providing important insights into the assembly and function of microbial communities across a range of ecosystems and indications including crop health, infectious disease, and autoimmunity. An important part of these experiments is understanding how abundant and prevalent the strains within the community are. A unique property of these synthetic communities is that they are normally constructed from genome-sequenced constituents which provides a constrained reference for the set of reads that may arise from metagenomic sequencing.

Traditional methods for quantifying organism abundances, such as 16S rRNA gene sequencing, usually lack the resolution to differentiate strains and are limited to generalizing to the species, genus, or higher taxonomic levels as a function of divergence within the clade of interest [11]. Much sequencing data, including shotgun metagenomic sequencing, is often quantified using the metric mapped fragments per kilobase per million reads (FPKM). The limitation of this approach for strain-resolved abundances is how to deal with reads that map to multiple strains, or ambiguous reads. Partially or randomly assigning ambiguous reads to all genomes that they map to introduces noise and inflates the abundance of low abundance and/or absent strains. Ignoring these reads, on the other hand, may lead to a bias where genomes that are more similar to each other have artificially reduced observed abundances. As such, NinjaMap [7] and StrainR [12], here referred to as StrainR1, were developed to address these challenges in a 119-member community of diverse gut microbes, and a 22-member community of entirely *Eggerthella lenta* strains [12], respectively. Both approaches explicitly include a normalization strategy to correct for the uniqueness of the target genome; though, they differ significantly in their implementations.

NinjaMap tackles the problem by using the proportion of uniquely mapped reads to partially assign ambiguously mapped reads to optimize use of all the reads available from a sequencing run. In addition, reads are generated *in silico* as part of the pipeline to assess and normalize for strain uniqueness. This does not necessarily solve the bias observed from uniquely mapped reads preferring more unique genomes, as the bias may be propagated to the assignment of ambiguously mapped reads. NinjaMap, therefore, exhibits some of the same problems as data which has not been normalized for unique mapping sites, as demonstrated in our analyses. Furthermore, this approach may lead to high false positive rates as it becomes likely that a genome has at some number of uniquely mapped reads through either sequencing error, trace cross contamination with input communities, or index hopping/barcode switching [13,14]. Another problem faced by NinjaMap is that it scales non-linearly, requiring extensive resources for complex communities and quickly running out of the resources required to run for larger communities. StrainR1 was developed to address the same issue. Unlike NinjaMap, it aims to only use uniquely mapped reads and normalize the bias towards more unique genomes directly by using the number of unique k-mers within each genome. Unfortunately, the implementation of StrainR1 fell short of being able to scale to larger and more diverse communities requiring excessive computational resources. This presents a need for a tool that can scale well enough that it can run in most computing environments for large synthetic communities while still maintaining the same, or better, accuracy.

Other tools for consideration include Strainer [15], which was designed to detect the presence or absence of strains implanted into an undefined community using fecal microbiota transplants (FMTs); however, it is not appropriate for use in the analysis of synthetic communities due to its intended application in undefined communities to find informative k-mers against a background community. Another alternative for determining presence or absence of strains which may be applied to synthetic communities is YACHT [16]: a tool that determines strain presence based on metagenomic sequencing reads. Neither of these approaches is designed for quantitative analysis of strains, but can still be applied to benchmark the presence/absence detection of StrainR2.

In this manuscript, we introduce and benchmark StrainR2 which quantifies strain abundances to a higher degree of accuracy than other methods relying on shotgun metagenomic sequencing data while maintaining scalable and lightweight run times and resource usage. StrainR2 can run both in a high-performance computing environment and on a personal computer. In our analyses, we find that the currently available tools derive inaccurate abundances, particularly for cases of low abundance organisms or communities that contain highly similar strains. We further validate the superior accuracy of StrainR2 by benchmarking against real samples which were characterized using strain-specific qPCR against synthetic communities colonizing gnotobiotic animals. StrainR2 will facilitate synthetic microbial community research of increasing complexity as it scales to study complex host-associated microbiomes like the mammalian gut.

## IMPLEMENTATION

StrainR2 has two steps: (i) preprocessing (PreProcessR) and (ii) normalization (StrainR, **Figure 1**). In short, the preprocessing step first splits every genome’s contigs into subcontigs such that each genome has a comparable build quality. The number of unique k-mers in each subcontig is then calculated as a measure of a strain’s uniqueness. This measure is used in the normalization step for part of a modified FPKM formula termed fragments per thousand <u>u</u>nique <u>k</u>-mers per <u>m</u>illion reads mapped (FUKM). A user-configurable weighted percentile of all subcontig FUKMs in a genome is used as the final measure of abundance. FUKM is directly analogous to FPKM, with the difference of normalizing the uniquely mappable sites of the genome rather than the total genome size.

**Figure 1.**
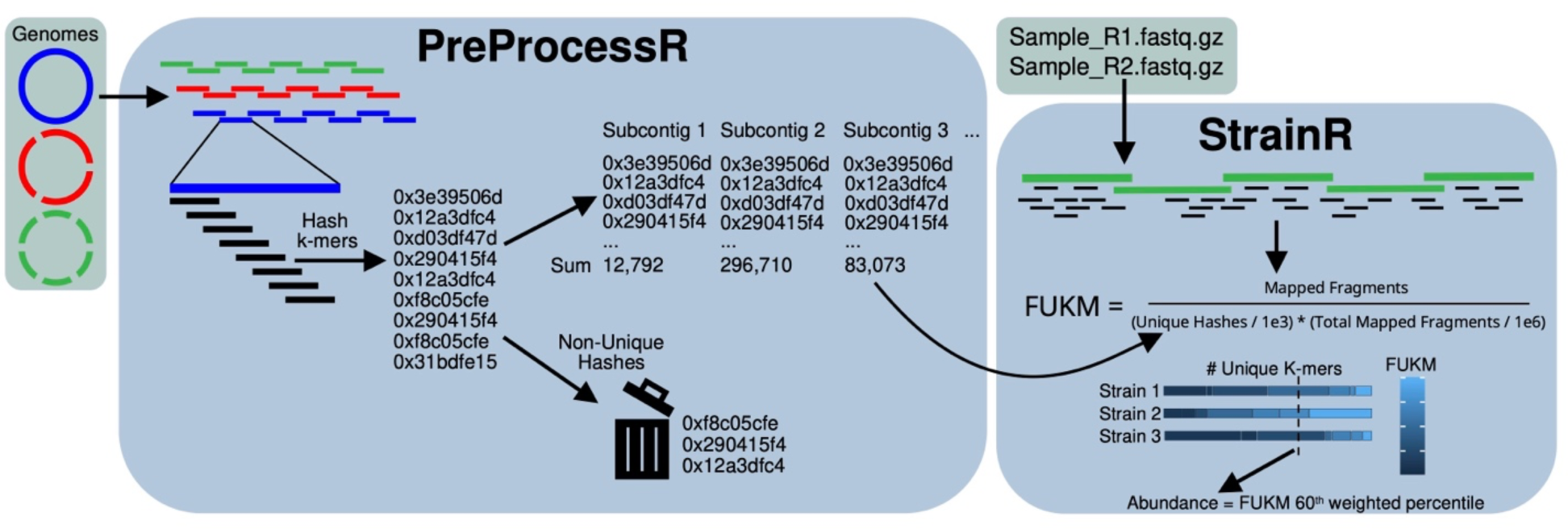
Schematic of the StrainR2 workflow. Genomes are split into subcontigs no larger than the smallest N50 in the set of genomes ensuring consistent assembly qualities. The number of unique k-mers (which are computed as hashes for efficiency) is used to normalize FPKM in a metric normalized for genome uniqueness (FUKM). A user-configurable weighted percentile of all subcontig FUKMs belonging to a genome is used as a point estimate of abundance.

### Preprocessing

The preprocessing step is almost entirely written in C to maximize efficiency. It starts by splitting contigs of all genomes into similarly sized pieces to ensure the build quality for all genomes is comparable. This eliminates bias towards build quality and also ensures there are sufficient subcontigs for use in the normalization step by providing multiple estimates of strain abundances and thereby normalizing out the effects of highly unique multi-copy elements such as plasmids and transposons. By default, the smallest N50 build quality for all provided genomes is used as the maximum possible size of subcontigs. StrainR2 maximizes subcontig size given the constraint of the smallest N50 being the upper bound. Subcontigs also have a 500 base overlap with the preceding and proceeding subcontigs to ensure reads in between subcontigs are mapped. Contigs under a minimum size, set to 10 kbp by default, are excluded from the final calculation for FUKM but will still have their k-mers marked as non-unique. These small contigs could represent multi-copy elements and therefore bias read mappings if their k-mers are considered unique.

Canonical k-mers are generated and represented as 64-bit hashes for the sake of computational speed. K-mers are always made to be odd so that no k-mer can be the same as its reverse complement. The non-cryptographic MurMurHash is used as the hashing function due to its speed. The default k-mer size matches the total number of bases in both paired-end reads with the logic that the number of unique k-mers is an approximation of the number of sites a read can uniquely map to. Unique hashes (or k-mers) in a subcontig are measured as being unique with respect to all genomes in the community. To determine which k-mers are unique in each subcontig, StrainR2 iteratively adds each k-mer’s hash to a hash table of all hashes encountered in the community thus far. As this is done, a count for the number of unique hashes in each subcontig is maintained and counts are only correct given the k-mer hashes processed so far. This means that if a k-mer hash is encountered that was previously considered unique, it is marked as non-unique and the unique k-mer count for its respective subcontig is decremented by one. The hash table is implemented with open addressing and a linear probe collision policy, as hash table entries are relatively small. The hash table is resized to powers of two whenever the load factor exceeds 0.75 after the hashes for a subcontig are added.

One caveat to using hashes in place of k-mers is the possibility that two k-mers may have the same hash (collision). Assuming 150 base paired-end-reads are used (leading to the default use of 301-mers), that indicates that 1.7e181 possible k-mers will map to 1.8e19 possible hashes. However, a typical synthetic community has on the order of 10s or 100s of millions of different k-mers, meaning the probability of different k-mers having the same hash is low and the effect is negligible.

The preprocessing step only needs to be performed once per community, with the normalization step using the values found in preprocessing to normalize each set of reads.

### Normalization

In the normalization step, reads are trimmed using fastp [17] with the options: trim_poly_g, length_required=50, n_base_limit=0. BBMap [18] is subsequently used to obtain the count of fragments that perfectly and unambiguously map to each subcontig, with the options: perfectmode=t, local=f, ambiguous=toss, pairedonly=t. Finally, FUKM is calculated using the formula: 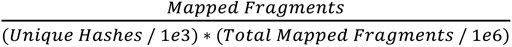. To obtain a singular value for the abundance of a strain, a user-configurable percentile of subcontig FUKMs is recommended. The percentiles are also weighted by the number of unique k-mers in the subcontig so that percentiles are more representative of areas of the genome that were more likely to be uniquely mapped to. The weighted percentile of FUKMs for each strain is abbreviated wpFUKM, and the median FUKM is abbreviated mFUKM.

### Unit testing

The stability and reproducibility of StrainR2 is validated by automated testing (GitHub workflow) to ensure that the output at each step of the workflow is reproducible and deterministic. Testing includes running PreProcessR and StrainR as shell commands in a conda environment, as well as running individual components of each command separately. For PreProcessR, this includes testing that genome contigs are correctly and completely divided into subcontigs, as well as ensuring that unique k-mer hash counts are correct. For StrainR, the calculation for FUKM is validated. This testing also ensures that StrainR2’s output remains consistent through updates to itself or its environment, as testing is triggered on any change to the source code.

### *In silico* read generation

For the purpose of assessing accuracy and speed, InSilicoSeq [19] was used to generate 20 million reads for varying distributions across two different communities. The coverage option was used with predetermined coverages to simulate these distributions **(Figure S1, Tables S1-4)**. Reads from a NovaSeq 6000 S4 flow cell, which generated later experimental validation data, were used to create a custom error model for all *in silico* reads.

### Tool analyses

Unless otherwise noted, StrainR2 was always run with default options: a weighted percentile of 60; max subcontig size of the lowest N50 in all input genomes; minimum subcontig size of 10Kb; read size of 150; no subcontig filtering. NinjaMap was also run with default options.

FPKM values were taken from the same mapping data that StrainR2 normalizes. More specifically, BBMap was run with the options: perfectmode=t, local=f, ambiguous=toss, pairedonly=t, and nodisk=t. All other options were left to default. The FPKM was then calculated by summing the mapped fragments for each contig belonging to a strain in the .rpkm file and normalizing it by kilobases and millions of total reads.

To test StrainR2’s ability to determine strain presence/absence, it was benchmarked against YACHT [16]. While it does not provide abundances, it can still be benchmarked against StrainR2 based on if StrainR2 provides an abundance of 0 or not. YACHT was run with k-mers of size 31 with the scaled option set to 100. Significance was set to 0.99 and a minimum coverage of 0.01 was used.

### Resource expenditure profiling

Tool preprocessing steps were tested on varying input sizes of random genomes belonging to human gut microbes from our in-house strain collection: between 10 and 200 genomes, as well as varying input sizes of the 22 *E. lenta* strains. Testing was done on an Ubuntu server with dual Intel Xeon Silver 4214 CPUs and 384 GiB of memory. When resource usage for any tool became too large to be run with the computing resources available, or if the tool threw an error, testing was stopped.

### Experimental validation

Mice (strain C57BL/6J) aged 8-17 weeks were given *ad libitum* Lab Diet 5021 and had a 12-hour light/dark cycle. Mice were housed inside Class Biologically Clean germ-free isolators in the gnotobiotic animal facility at Pennsylvania State University. Synthetic communities were assembled by pooling approximately equal cell densities of individual strains described elsewhere (**Table S1**)[9]. Mouse fecal samples were extracted following the International Human Microbiome Consortium Protocol Q [20]. Briefly, samples were homogenized through bead disruption before isopropanol precipitation and extraction using the Qiagen Stool DNA kit. Libraries were prepared using the Illumina Library Preparation kit and analyzed using a NovaSeq 6000 (Novogene USA). Data is available for download at PRJNA1038784. Negative controls and fecal material from germ-free animals were also prepared and pooled for sequencing despite failing library construction QC. Reads from these communities were mapped primarily to the mouse genome and/or common reagent contaminants.

Strain abundances were quantified using qPCR for JEB00023, JEB00029, JEB00174, and JEB00254 (**Table S1**). Primers were designed using PrimerBLAST to be specific to unique portions of each target genome. Primers were validated *in silico* and experimentally using gDNA from the pure culture members of the sFMT1+Cs community via qPCR to ensure a lack of cross reactivity. The primers used are as follows: JEB00023 forward primer: GGCACTCATCGGAGGTTTCA, JEB00023 reverse primer: CGTTGGGCTTGTCACCAAAG JEB00029 forward primer: TCATGGCCGTGTACTTGCTT, JEB00029 reverse primer: AGCGGATATCTGCCAGGTTG, JEB00174 forward primer: TGGAGTTCGGCGTAGCTTTT, JEB00174 reverse primer: TCTCGGCATTCCAACCAGAC JEB00254 forward primer: ACAGGCTTTGGCATTGGAGA, and JEB00254 reverse primer TGTGGTTAATGGCCTTGCAT, with the concentration of each being 200 nM. Samples were amplified with iTaq Universal SYBR Green Supermix (Biorad 1725122) at 95°C for 3 minutes, followed by 40 cycles of 5 seconds at 95°C and 30 seconds at 60°C using a Biorad CFX384 Opus. Strains were quantified against standard curves of pure gDNA and normalized by extracted sample mass to calculate absolute abundance.

## RESULTS AND DISCUSSION

Throughout analysis, two communities of strains were used to generate reads *in silico* across 6 abundance distributions that represent various scenarios **(Figure S1, Tables S1-S4)**. The first community, sFMT1+Cs, represents a more realistic scenario for a synthetic community where most strains are not very similar but with five of the species having two or more strains each **(Figure 2A, Table S1)**. As in the development of StrainR1, a mock community of 22 *E. lenta* strains were also used to represent an extreme scenario where the number of unique k-mers would be at a minimum **(Figure 2B)**.

**Figure 2.**
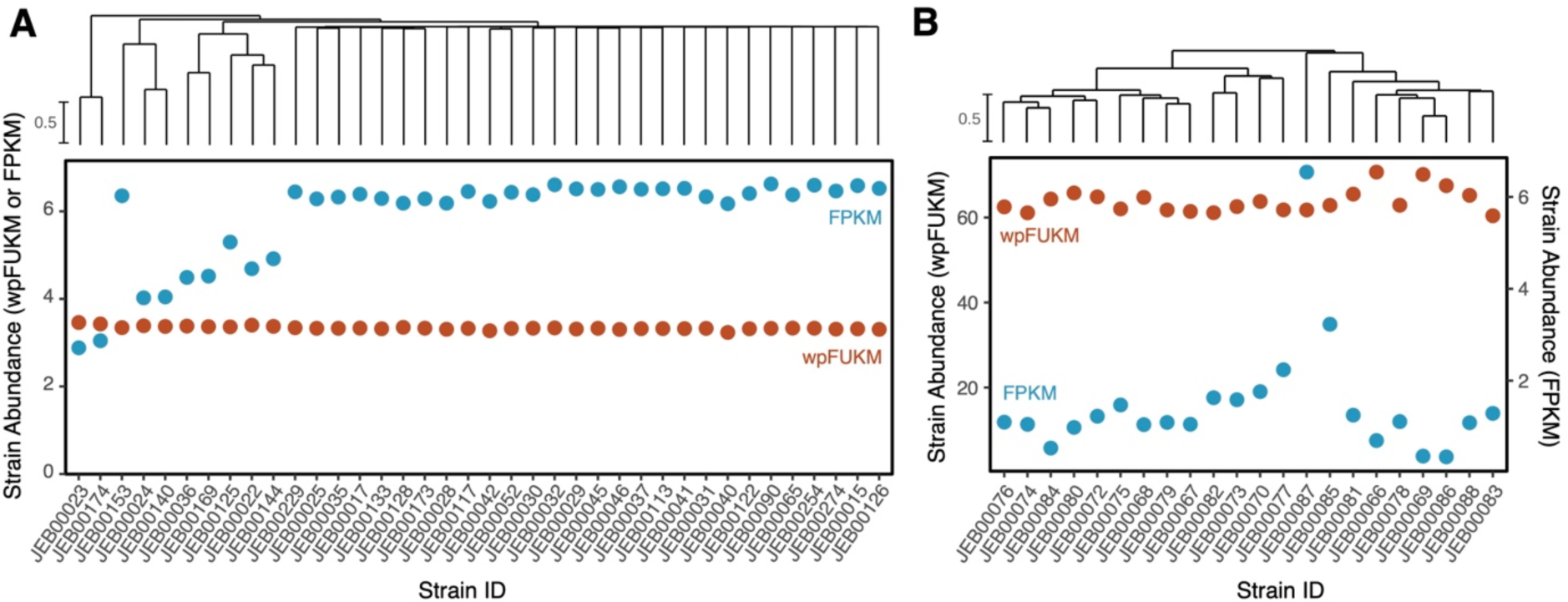
StrainR2 normalization corrects quantitative errors resulting from variable strain-relatedness. Dendrograms of strain similarities are shown for **(A)** sFMT1+Cs members and **(B)** 22 *E. lenta* strains. Reads were generated *in silico* such that all community members have a uniform abundance. StrainR2 resolves abundances much closer to the uniform abundance than by measuring FPKM. FPKM values for *E. lenta* strains were less accurate than with the sFMT1+Cs community due to an increased bias towards unique community members. StrainR2-calculated wpFUKM had a coefficient of variation of 1.69% for the sFMT1+Cs strains and 3.93% for the *E. lenta* strains despite the high strain similarity, whereas when using FPKM the coefficient of variation was 17.44% and 86.82% for the communities, respectively. Dendrograms are based on Jaccard Similarity of k-mer profiles between strains.

To evaluate StrainR2’s improvement over using FPKM values in the case of uniformly abundant strains, the coefficient of variation was used as a benchmark, as lower coefficients of variation are closer to a uniform distribution. In the case of the sFMT1+Cs community with uniformly abundant strains, StrainR2’s wpFUKM was able to normalize the reads such that the coefficient of variation was 1.69% as compared to FPKM with a coefficient of variation of 17.44% **(Figure 2A)**. In the case of uniformly abundant *E. lenta* strains, the coefficients of variation were 3.93% and 86.82% for wpFUKM and FPKM, respectively **(Figure 2B)**. Using StrainR2, this corresponds to fold change differences of 1.085 and 1.159 between the highest and lowest reported abundances for wpFUKM on sFMT1+Cs and *E. lenta* strains, respectively. Despite the high similarity between strains, wpFUKM still resembled a uniform distribution unlike FPKM. FPKM tended to underestimate the abundance of strains with higher similarity, whereas wpFUKM remained unbiased.

With uniformly abundant *E. lenta* reads, StrainR2’s wpFUKM best followed a uniform distribution out of all methods tested as determined by coefficients of variation **(Figure S2A)**. While median FUKM (mFUKM) is the only abundance estimate provided by StrainR1, StrainR2 is still able to quantitatively improve on this measure, with the coefficient of variation decreasing from 6.22% to 5.53%. Specifically, StrainR2 achieves this by including overlaps between subcontigs, using larger k-mers, and marking k-mers from excluded subcontigs as non-unique. Furthermore, NinjaMap showed the worst performance out of all the methods tested, with several cases of strain abundances being off by more than tenfold **(Figure S2B)**.

To further assess the accuracy examined the recovery of strain abundances across 6 different distributions (**Figure S1, Tables S3 and S4**). Jensen-Shannon divergence between true and predicted abundances was used. All measures of abundance were first converted to be a percentage of the total abundance so that all measures of abundance were comparable, then the Jensen-Shannon divergence was calculated. Resulting values can be between 0 and 1, where 0 represents the least divergence. Across all types of distributions, StrainR2 had a Jensen-Shannon divergence at least two magnitudes smaller than either NinjaMap or FPKM **(Figure 3A)**.

**Figure 3.**
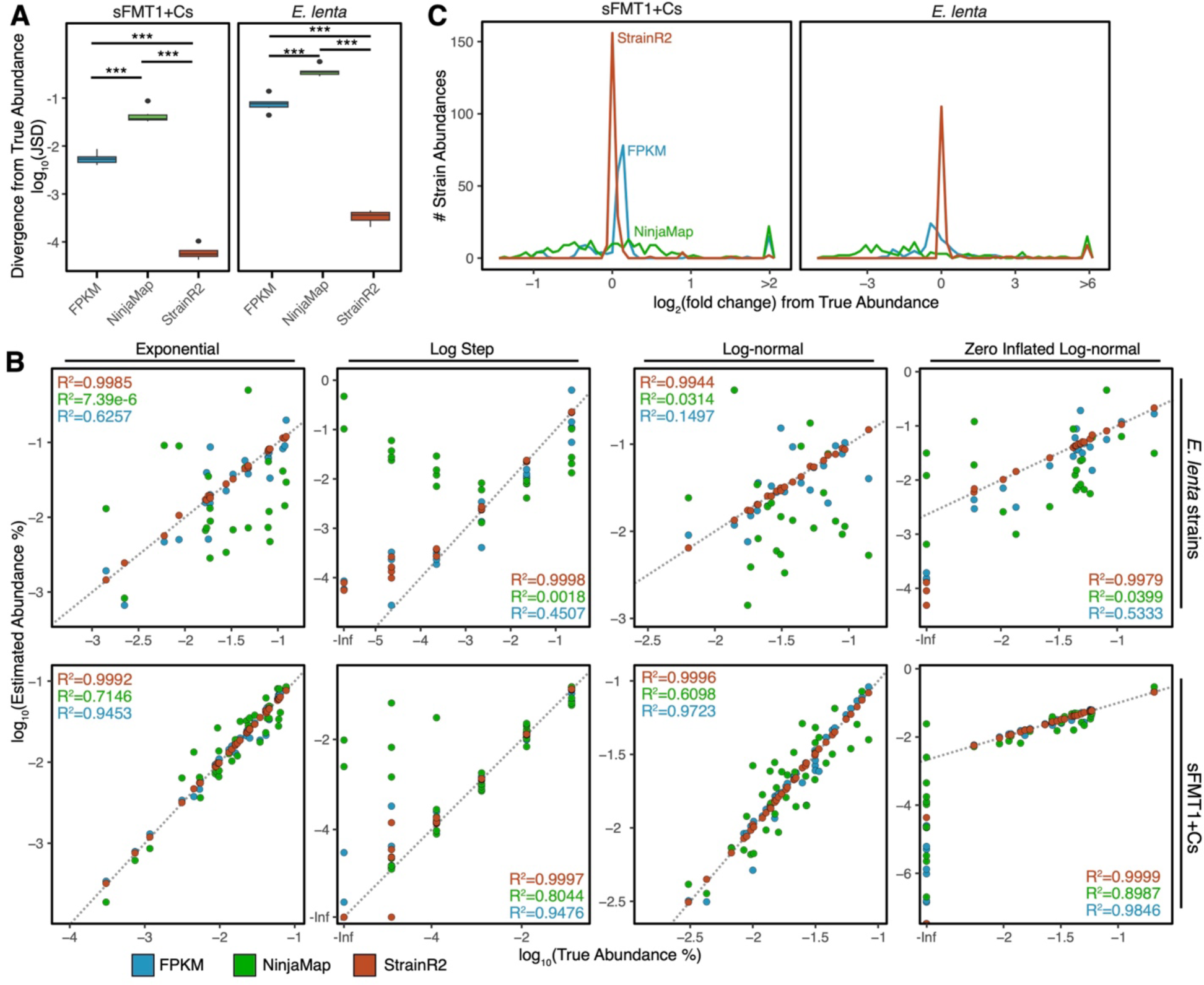
StrainR2 provides accurate strain abundances across varied community compositions. **(A)** Jensen-Shannon divergence between true community composition and estimated abundance are much closer to 0 than when using NinjaMap or FPKM. *** denotes a significance of less than 0.001, as determined by ANOVA and TukeyHSD. Reads were generated *in silico* across multiple community compositions and distributions **(Figure S1)**. **(B)** Scatterplots for the correlation between estimated abundances and true community compositions are shown for all mock community distributions in sFMT1+Cs and *E. lenta* strains. Inset values represent the Pearson correlation for each tool. The uniform and missing distributions are omitted as most or all strains have the same true abundance render the scatterplots non-informative. **(C)** A frequency plot of the fold change from the true abundance shows that StrainR2 rarely predicts an abundance far from the true abundance. Data shown is the sum of all 6 mock community distributions both for sFMT1+Cs and *E. lenta* strains.

Across each distribution, StrainR2 consistently provided the most accurate recovery of relative abundances (**Figure 3B**). NinjaMap’s abundance predictions were less accurate than using FPKM in all community distributions, which results from high abundance predictions for the strains with the lowest abundances. Its accuracy degraded significantly when spanning strains with orders of magnitude difference in abundance. Moreover, the *E. lenta* community decreased the accuracy of all tools, but StrainR2 observed the smallest decrease in accuracy. As a representative example, StrainR2 more closely correlates with true abundances in a log-normal distribution of sFMT1+Cs strains than other methods. StrainR2 had the highest Pearson correlation, with an R^2^=0.9996, whereas NinjaMap and unnormalized FPKM reported correlations of R^2^=0.6098 and R^2^=0.9723, respectively. In the log-normal distribution of *E. lenta* strains, the R^2^ values are decreased to 0.9944 (StrainR2), 0.1497 (FPKM), and 0.0314 (NinjaMap). The scatterplots for other distributions and communities show a similar trend with NinjaMap and FPKM performing far worse with *E. lenta* strains, whereas StrainR2 maintains high correlations (R^2^>0.9944) **(Figure 3B)**.

The frequency of log2(fold changes) from the true abundance summed across all 6 distributions for each tool shows that StrainR2 has the largest peak around 0 with few abundance predictions more than two fold different from the true abundance **(Figure 3C)**. NinjaMap had the largest amount of predictions at more than a 4 fold change from the true abundance, which mostly arises from low abundance organisms. These could be a result of NinjaMap’s use of reads that map ambiguously, which inflates abundances in the case where most ambiguously mapped reads originate from another organism. StrainR2 has similar such cases for very low abundance strains, but at a significantly reduced rate.

StrainR2 can also be used to test the presence or absence of a strain depending on if it outputs zero as an abundance. The rate of false positives and negatives heavily depends on which weighted percentile is used as the final abundance **(Figure S3A)**. Using a higher weighted percentile increases false positives, whereas lower weighted percentiles increase false negatives, usually in the case of low abundance organisms. A weighted percentile of 60 was chosen to compare StrainR2’s strain presence or absence predictive ability. F1 scores of StrainR2 and three other tools reveal that StrainR2 predicted presence with the highest accuracy **(Figure S3B)**. Only three distributions are shown as these are the only distributions that contained absent strains. These results suggest that StrainR2 performs slightly better than YACHT for predicting the presence or absence of strains with recommended parameters, and shows a large improvement over NinjaMap and FPKM. To further test the presence/absence prediction of these tools, the tools were run on ten replicates of the zero-inflated log-normal community, each with different strain abundances, with StrainR2 still showing the best performance **(Figure S3C)**.

To assess how StrainR2 scales with synthetic community complexity, the memory in GB and run times of the computationally intensive database generation steps of StrainR2, StrainR1, and NinjaMap were gathered as described in the implementation section on an Ubuntu server with dual Intel Xeon Silver 4214 CPUs and 384 GiB of memory (**Figure 4**). Run times for StrainR2 grew linearly and remained low for all inputs as compared to StrainR1 **(Figure 4A)** and the memory usage showed similar trends **(Figure 4C)**. To test if the trend would hold for highly similar inputs, runs were performed on one through 22 of the *E. lenta* strains. StrainR2 maintained its low run times, whereas StrainR1 and Ninjamap’s run times grew non-linearly **(Figure 4B)**. Memory usage on the *E. lenta* strains is also shown **(Figure 4D)**. StrainR2 run times scale closely with the number of unique k-mers there are in a community, meaning it is unaffected by highly similar communities **(Figure S4)**. To validate function outside of a high-performance computing environment, the sFMT1+Cs community was profiled through a StrainR2 Bioconda installation on a personal computer. The system was running OS X Ventura 13.1 with an Apple M1 Pro processor and 16Gb of RAM. The run times were 1 minute 31 seconds, and 6 minutes 56 seconds for PreProcessR and StrainR respectively for 51.9 million reads (7.8 Gbases) of paired-end NovaSeq 6000 data with 8 threads. This highlights that StrainR2 does not require high performance computing nodes, and is a tangible strategy available to most research groups.

**Figure 4.**
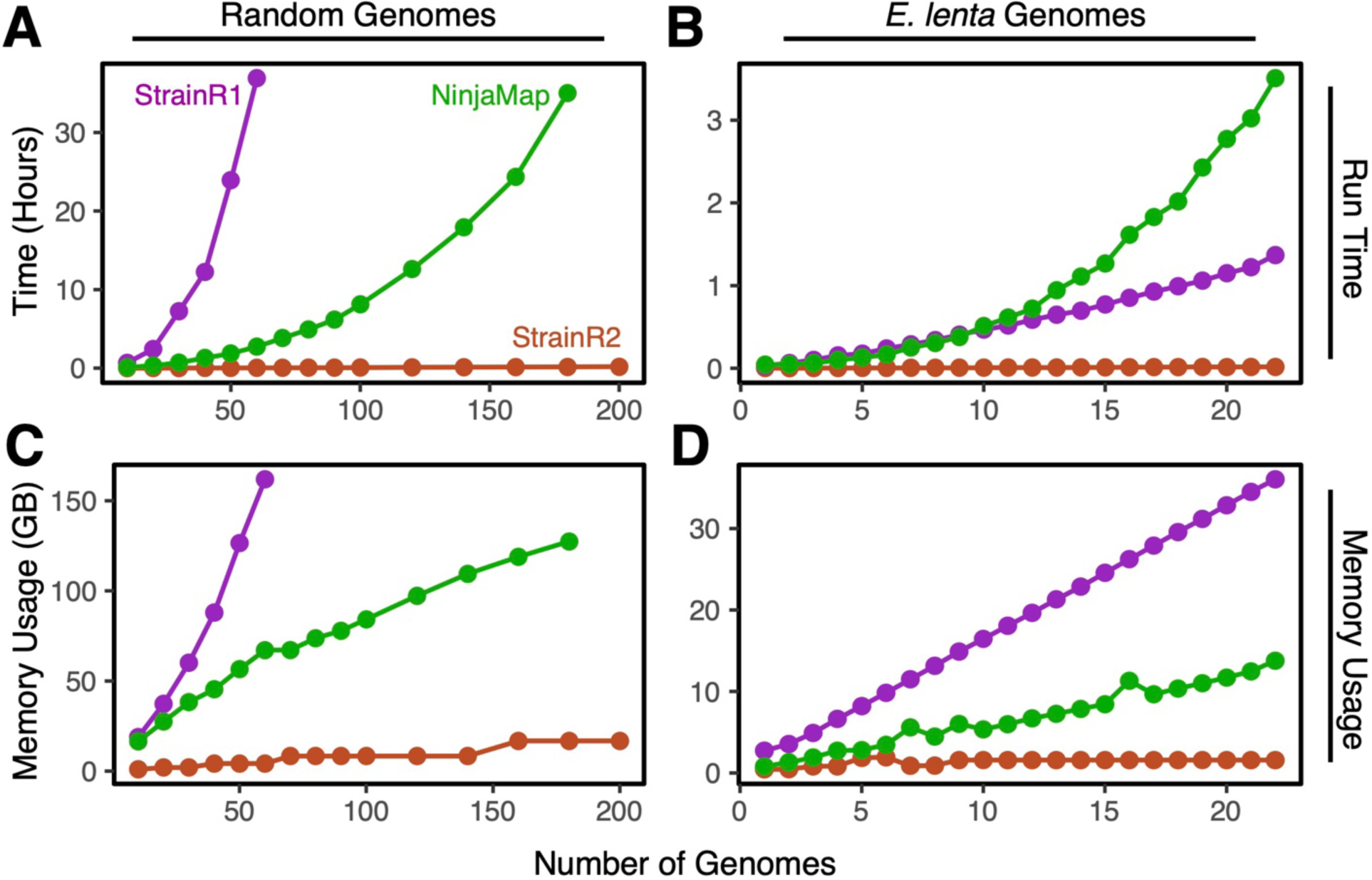
StrainR2 uses fewer system resources while scaling linearly. Run times for database generation for **(A)** 200 random genomes and **(B)** *E. lenta* strains show StrainR2 following a linear growth. StrainR2’s final run times were 3 minutes and 50 seconds, and 32 seconds, respectively at maximum community complexity for random genomes and *E. lenta* respectively. Memory usage for **(C)** the 200 genome input and **(D)** the *E. lenta* community again shows StrainR2 using the least resources, with final memory usages of 16.8 GB and 1.4 GB, respectively.

While StrainR2 was shown to have the best accuracy *in silico*, we sought to validate its function using experimental samples and a gold-standard method for strain quantification: qPCR with strain-specific probes. Shotgun metagenomic sequencing data was obtained from 17 fecal samples of gnotobiotic mice colonized with sFMT1 or sFMT1+Cs. As a control, two of the mice were also germ-free. To determine the true abundance of strains, qPCR was performed on samples from the same mice for four of the strains present in sFMT1 (JEB00023, JEB00029, JEB00174, and JEB00254). JEB00023 and JEB00174 were selected as they are both strains of the species *Bacteroides uniformis* and represent an important use case of StrainR2. JEB00029 was selected as our previous experiments had suggested it could not colonize the mice, while JEB00254 was capable of colonization at low abundances [9]. To compare abundances from each method, the fold change from the geometric mean of strain abundances within a sample was calculated.

StrainR2 maintained a close relationship with the results obtained from qPCR, as well as correctly predicting when strains were absent, as was the case with JEB00029 and the two germ-free mice **(Figure 5A)**. NinjaMap showed a weaker correlation with the data from qPCR, and was inconsistent with predicting abundances of strains between samples **(Figure 5B)**, with FPKM having similar results **(Figure 5C)**. FPKM and NinjaMap both incorrectly assigned abundances to JEB00029 and strains in the germ-free mice, showing that another strength of StrainR2 is more accurate presence/absence prediction, as it was the only tool to agree with qPCR on the absence of strains. Per-strain correlations with each tool are described in **Figure 5D**. Pearson correlation values for StrainR2, NinjaMap, and FPKM versus copies/g across all strains are R^2^=0.9432, R^2^=0.3139, and R^2^=0.3559, respectively.

**Figure 5.**
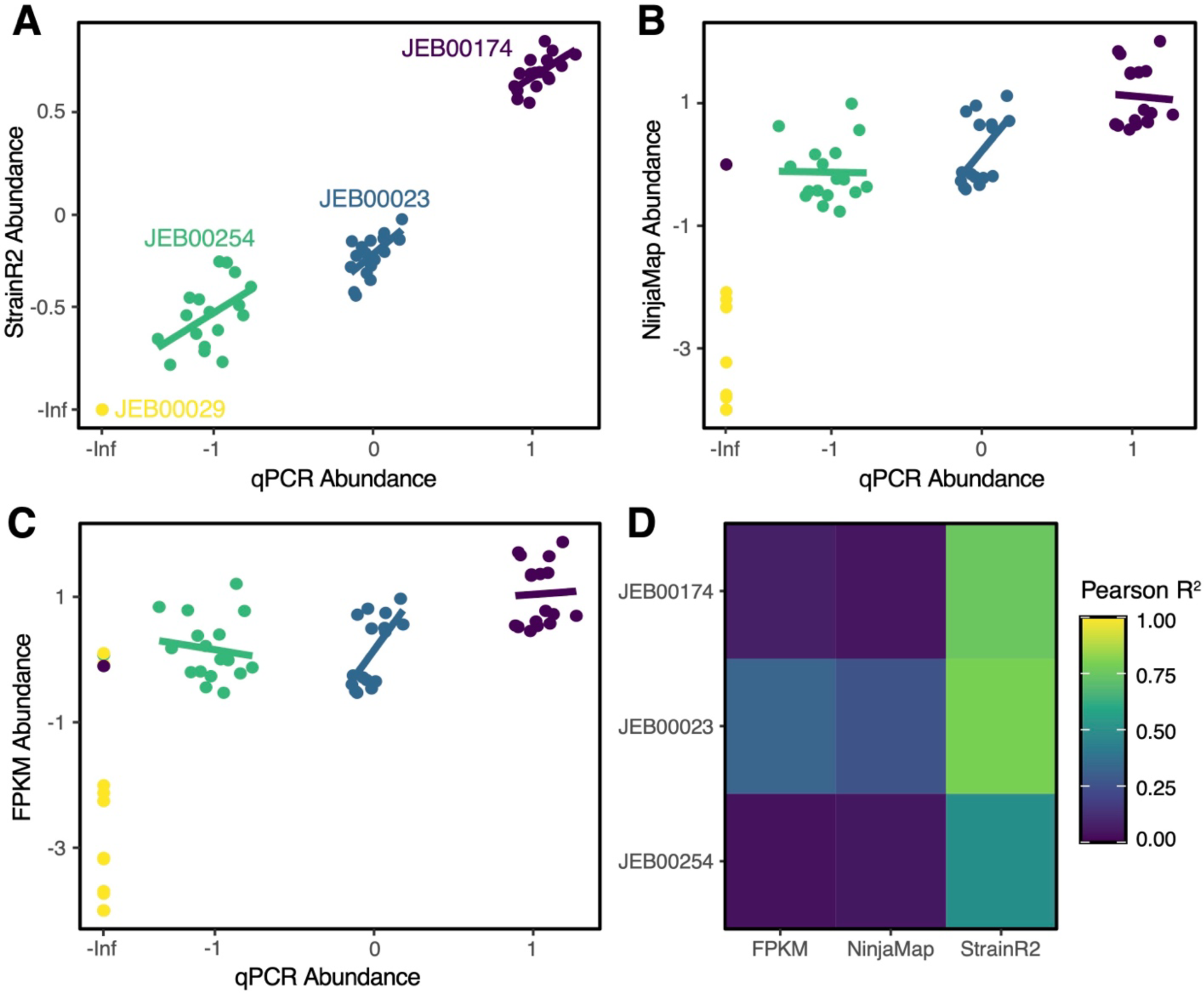
StrainR2 accurately recovers abundances measured by qPCR. **(A)** StrainR2 predicts abundances between samples and strains with the highest degree of accuracy as compared to **(B)** NinjaMap and **(C)** FPKM. **(D)** StrainR2’s predicted abundances are most closely correlated with the absolute abundance predicted by qPCR. It was also the only tool to correctly predict the absence of JEB00029 in all cases. All abundances are scaled as log_10_(fold change) from the geometric mean of strain abundances. Pearson correlations for all samples/strains in panels A, B, and C are R^2^=0.9432, R^2^=0.3139, and R^2^=0.3559, respectively. Each point represents the quantification of a single strain in a single animal with linear regressions drawn on a per-strain basis.

## CONCLUSIONS

Through analysis of data both *in silico* and experimental data, we demonstrate that StrainR2 provides highly accurate strain abundances and prevalences using a fraction of the computational resources of previous approaches. StrainR2 makes shotgun metagenomic sequencing reads a viable tool for accurate strain abundances in synthetic communities without the need for high performance computing. This may eliminate the need for more time consuming or expensive methods to assess strain abundance, such as qPCR to which it provides comparable abundances. StrainR2 is also able to provide abundances in scenarios where designing primers for qPCR would be extremely difficult or impossible, as would be the case for the *E. lenta* community. StrainR2 is available via GitHub, Bioconda, and as a Docker container.

## AVAILABILITY AND REQUIREMENTS

Project name: StrainR2

Project home page: https://github.com/BisanzLab/StrainR2

Operating system: Unix-like

Programming languages: C, Bash, R

Other requirements: conda (version >= 22.9.0)

License: MIT License

Any restrictions to use by non-academics: See license

## LIST OF ABBREVIATIONS

qPCR: Quantitative polymerase chain reaction
FPKM: Fragments per thousand bases per million reads
FUKM: Fragments per thousand unique k-mers per million reads
wpFUKM: Weighted percentile FUKM
mFUKM: Median FUKM
FMT: Fecal microbiota transplant

## DECLARATIONS

### Ethics approval

Animal experiments were approved by the Pennsylvania State University Institutional Animal Care and Use Committee.

### Availability of data and material

Gnotobiotic sequencing data is available via the NCBI Sequence read archive under BioProject PRJNA1038784. Source code and required resources to regenerate in silico reads are available at github.com/kheber/strainr2_mock_read_generation.

### Competing interests

The authors declare that they have no competing interests.

### Funding

Funding was provided by the NIAID (R00 AI147165) and NIGMS (R35 GM151045). Funding sources had no role in the design of the study and data collection, analysis, and interpretation of data.

### Authors’ contributions

K.H.: Conceptualization, methodology, software, formal analysis, writing - original draft, visualization. S.T.: Validation, investigation, writing - review & editing. D.B.- A.: Validation, investigation, writing - review & editing. H.K.: Validation, investigation, writing - review & editing. J.E.B.: Conceptualization, methodology, software, formal analysis, writing - original draft, visualization, supervision, funding acquisition.

## Acknowledgements

The authors wish to thank David Koslicki for helpful discussions.

## Supplemental Materials

**Figure S1.**
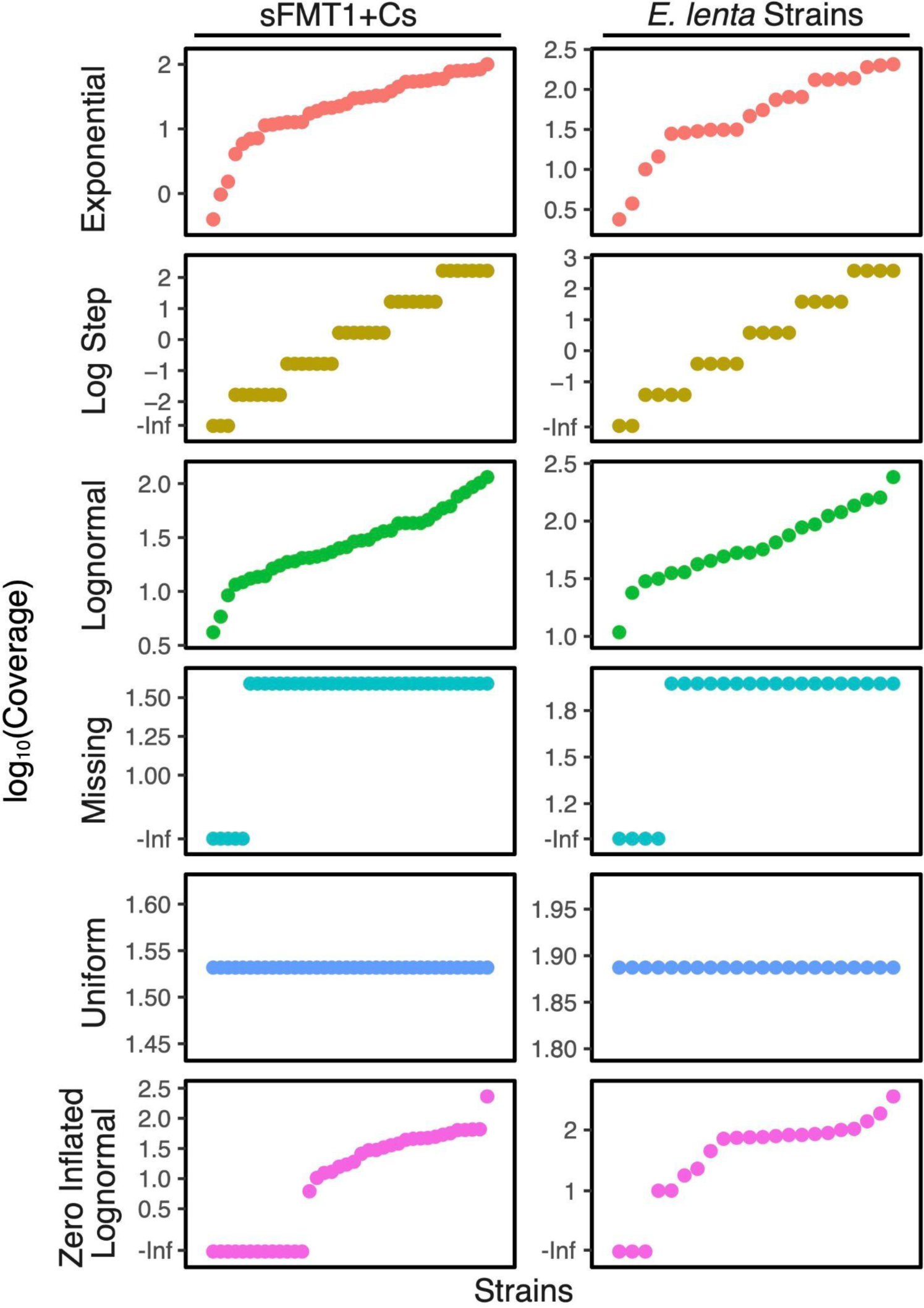
Mock reads simulate a variety of distributions. The 6 predefined *in silico* read distributions were designed to test the performance of tools in predicting presence/absence and scenarios where strains had low abundances or were not present. Abundances are defined as the read coverage depth of each strain. Strain coverages are reported in **Tables S3 and S4.**

**Figure S2.**
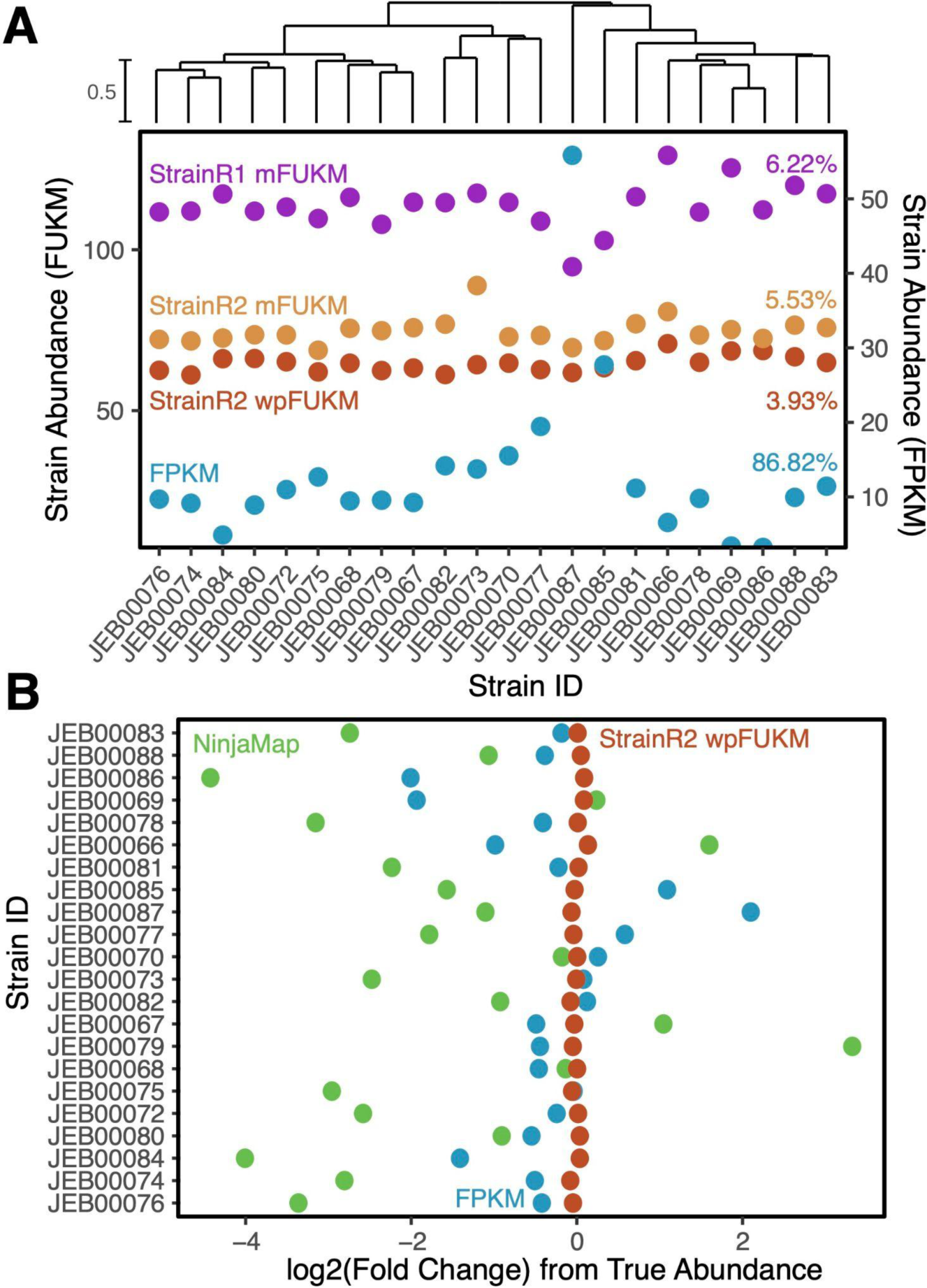
StrainR2 improves upon abundance estimation of *E. lenta* strains when compared to StrainR1. **(A)** Estimated abundances for a uniformly abundant community of *E. lenta* strains shows that StrainR2’s wpFUKM most closely follows a uniform distribution. Percentages shown to the right indicate coefficient of variation. **(B)** Accuracy as measured by log_2_(fold-change) demonstrates that StrainR2 is able to correct abundances to a high degree of accuracy, whereas NinjaMap and FPKM perform with much lower accuracy.

**Figure S3.**
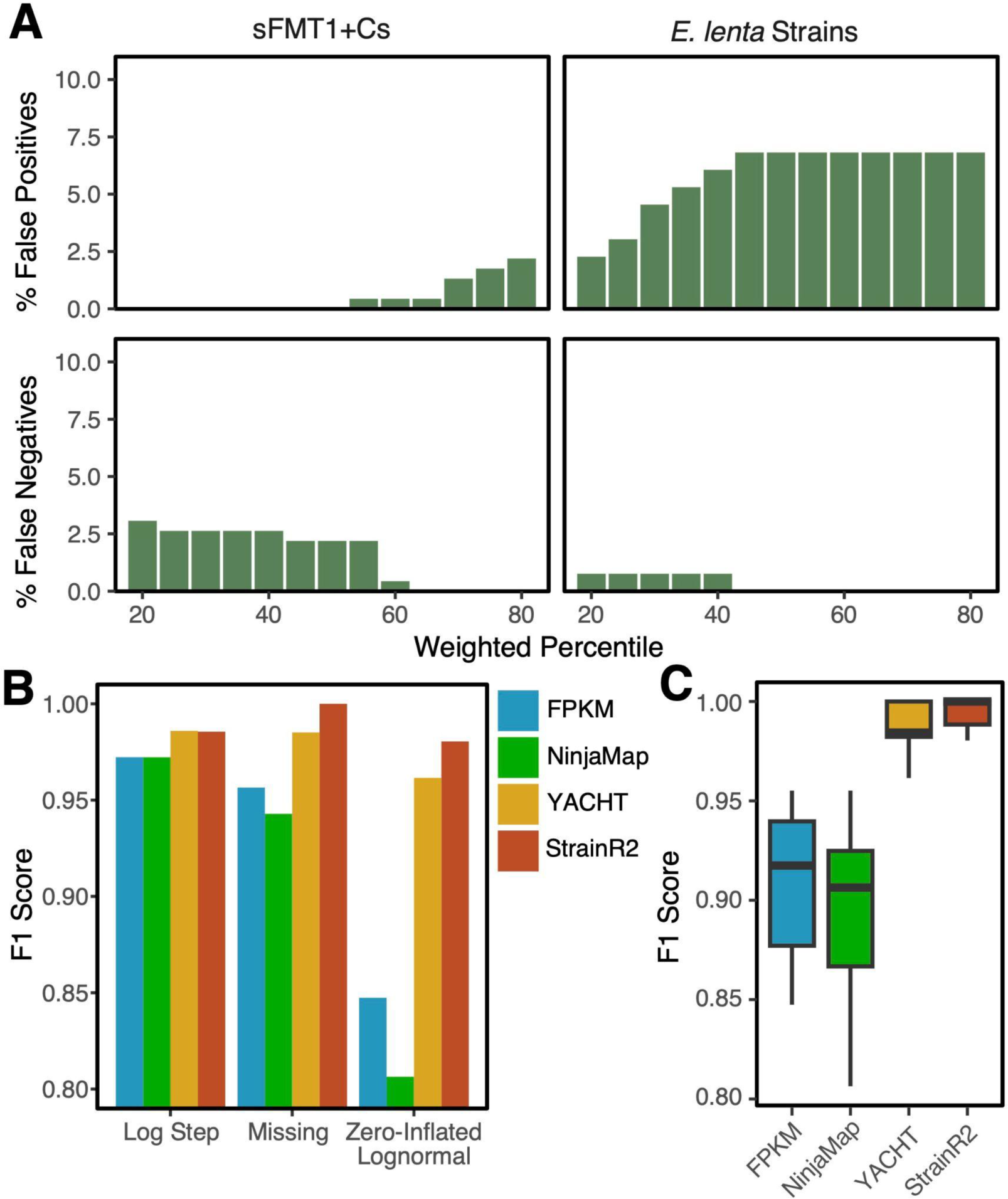
StrainR2 predicts presence or absence of strains with improved accuracy compared to other tools. **(A)** The average percentage of false positives or negatives across the 6 read distributions of sFMT1+Cs and *E. lenta* across different weighted percentiles of FUKM. **(B)** StrainR2 has the highest F1 scores across 3 of the read distributions and for the four tools. **(C)** Across ten replicates of the zero-inflated log-normal distribution, StrainR2 consistently has improved presence/absence prediction.

**Figure S4.**
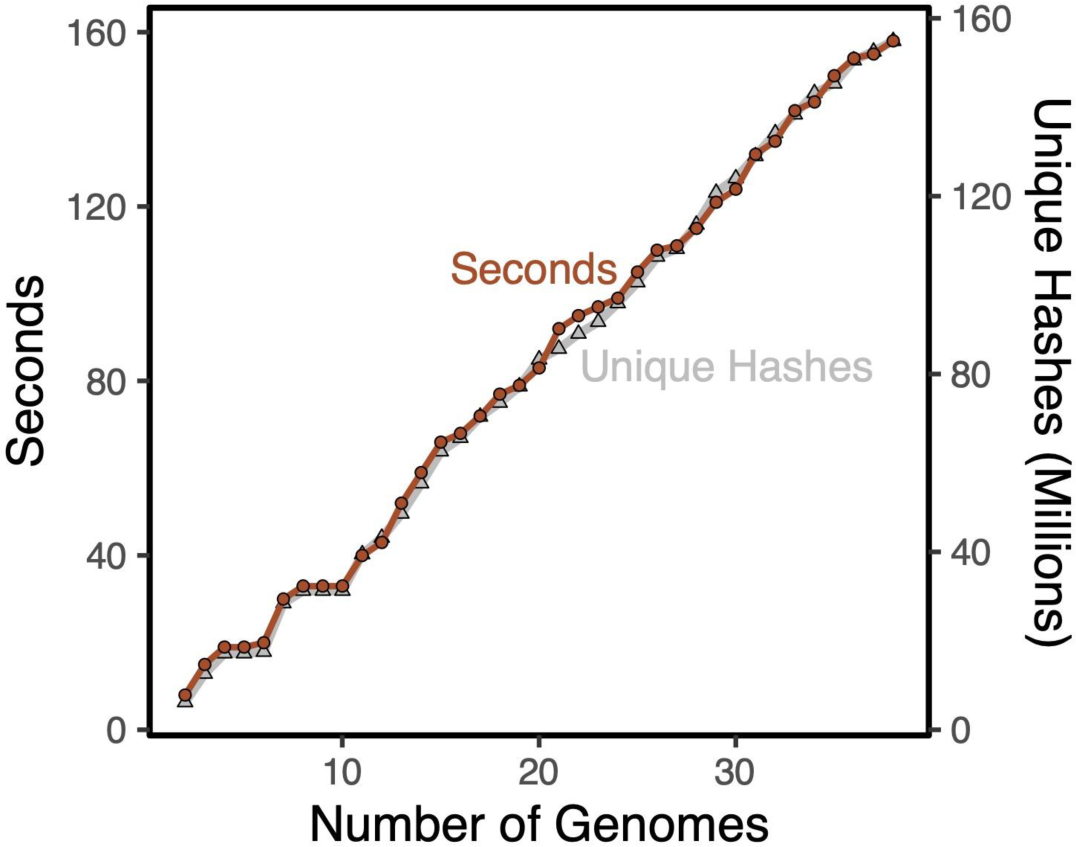
StrainR2 run times scale closely with the number of unique hashes in the input. Run times are plotted on varying input sizes from genomes in the sFMT1+Cs community. Total unique hash counts are shown, which estimate how many unique k-mers there are in the inputted genomes. Run times scale closely with unique hashes, meaning the more similar a community is, the better the run time will scale.

**Table S1.** Genome accessions for sFMT strains.

| LabID | Species | Strain ID | Genome Accession |
| --- | --- | --- | --- |
| JEB00015 | *Escherichia coli* | DSM 18039 | GCA_000005845.2 |
| JEB00017 | *Bacteroides thetaiotaomicron* | DSM 2079 | GCA_014131755.1 |
| JEB00022 | *Bacteroides ovatus* | DSM 1896 | GCA_001314995.1 |
| JEB00023 | *Bacteroides uniformis* | DSM 6597 | JBEUMQ000000000 |
| JEB00024 | *Bacteroides vulgatus* | DSM 1447 | GCA_000012825.1 |
| JEB00025 | *Parabacteroides merdae* | DSM 19495 | GCA_900445495.1 |
| JEB00028 | *Enterocloster asparagiformis* | DSM 15981 | GCA_000158075.1 |
| JEB00029 | *Dorea longicatena* | DSM 13814 | GCA_000154065.1 |
| JEB00030 | *Agathobacter rectalis* | DSM 17629 | GCA_000209935.1 |
| JEB00031 | *Clostridium scindens* | DSM 5676 | GCA_004295125.1 |
| JEB00032 | *Lachnospira eligens* | DSM 3376 | GCA_000146185.1 |
| JEB00035 | *Bacteroides stercoris* | DSM 19555 | GCA_900106605.1 |
| JEB00036 | *Bacteroides xylanisolvens* | DSM 18836 | GCA_000210075.1 |
| JEB00037 | *Anaerobutyricum hallii* | DSM 3353 | GCA_000173975.1 |
| JEB00040 | *Lactonifactor longoviformis* | DSM 17459 | GCA_002915525.1 |
| JEB00041 | *Faecalibacterium prausnitzii* | DSM 17677 | GCA_010509575.1 |
| JEB00042 | *Blautia producta* | DSM 3507 | GCA_002915535.1 |
| JEB00045 | *Dorea formicigenerans* | DSM 3992 | GCA_000169235.1 |
| JEB00046 | *Blautia obeum* | DSM 25238 | GCA_000153905.1 |
| JEB00052 | *Blautia producta* | DSM 2950 | GCA_010669205.1 |
| JEB00065 | *Clostridium spiroforme* | DSM 1552 | GCA_000154805.1 |
| JEB00090 | *Eggerthella lenta* | DSM 2243 | GCA_003339945.1 |
| JEB00113 | *Eubacterium hadrus* | DSM 3319 | GCA_000332875.2 |
| JEB00117 | *Clostridium orbiscindens* | NCBI 1_3_50AFAAA | GCA_000760655.1 |
| JEB00122 | *Clostridium symbiosum* | NCBI WAL-14673 | GCA_000189615.1 |
| JEB00125 | *Bacteroides ovatus* | NCBI D2 | GCA_000159075.2 |
| JEB00126 | *Bifidobacterium longum* | NCBI 35B | GCA_000261225.1 |
| JEB00128 | *Bacteroides caccae* | NCBI CL03T12C61 | GCA_000273725.1 |
| JEB00133 | *Bacteroides cellulosilyticus* | NCBI CL02T12C19 | GCA_000273015.1 |
| JEB00140 | *Bacteroides vulgatus* | NCBI CL09T03C04 | GCA_000273295.1 |
| JEB00144 | *Bacteroides ovatus* | NCBI 3_8_47FAA | GCA_000218325.1 |
| JEB00153 | *Bacteroides dorei* | NCBI CL03T12C01 | GCA_000273075.1 |
| JEB00169 | *Bacteroides xylanisolvens* | 2_1_22 | GCA_000162155.1 |
| JEB00173 | *Bacteroides finegoldii* | NCBI CL09T03C10 | GCA_000304195.1 |
| JEB00174 | *Bacteroides uniformis* | 4_1_36 | GCA_000185585.1 |
| JEB00229 | *Parabacteroides sp.* | D13 | GCA_000162275.1 |
| JEB00254 | *Peptostreptococcus anaerobius* | CC14N | SAMN42012254 |
| JEB00274 | *Sutterella wadsworthensis* | NCBI HGA0223 | GCA_000411515.1 |

**Table S2.** Genome accessions for *E. lenta* strains.

| **Lab ID** | **Strain** | **GenBank Accession** |
| --- | --- | --- |
| JEB00066 | Eggerthella lenta 1356FAA | GCA_000185625.1 |
| JEB00067 | Eggerthella lenta 11C | GCA_003340245.1 |
| JEB00068 | Eggerthella lenta 14A | GCA_003340255.1 |
| JEB00069 | Eggerthella lenta 22C | GCA_003340195.1 |
| JEB00070 | Eggerthella lenta 28B | GCA_003340165.1 |
| JEB00072 | Eggerthella lenta DSM11767 | GCA_003340045.1 |
| JEB00073 | Eggerthella lenta DSM11863 | GCA_003340015.1 |
| JEB00074 | Eggerthella lenta DSM15644 | GCA_003340005.1 |
| JEB00075 | Eggerthella lenta UCSF2243 | GCA_003339975.1 |
| JEB00076 | Eggerthella lenta Valencia | GCA_003339885.1 |
| JEB00077 | Eggerthella lenta AN51LG | GCA_003340155.1 |
| JEB00078 | Eggerthella lenta MR1n12 | GCA_003339935.1 |
| JEB00079 | Eggerthella lenta 1160AFAAUCSF | GCA_003340395.1 |
| JEB00080 | Eggerthella lenta 326I6NA | GCA_003340465.1 |
| JEB00081 | Eggerthella lenta AB8n2 | GCA_003340145.1 |
| JEB00082 | Eggerthella lenta AB12n2 | GCA_003340445.1 |
| JEB00083 | Eggerthella lenta CC82BHI2 | GCA_003340075.1 |
| JEB00084 | Eggerthella lenta RC46F | GCA_003339915.1 |
| JEB00085 | Eggerthella lenta CC75D52 | GCA_003340405.1 |
| JEB00086 | Eggerthella lenta CC86D54 | GCA_003340065.1 |
| JEB00087 | Eggerthella lenta W1BHI6 | GCA_003339875.1 |
| JEB00088 | Eggerthella lenta A2 | GCA_003340125.1 |

**Table S3.** Strain coverages in mock sFMT distributions.

| StrainID | Exponential | Log step | Log-normal | Missing Strain | Zero inflated log-normal | Uniform |
| --- | --- | --- | --- | --- | --- | --- |
| JEB00015 | 32.73 | 0.00 | 9.21 | 39.00 | 63.99 | 34.03 |
| JEB00017 | 4.09 | 0.00 | 29.03 | 0.00 | 0.00 | 34.03 |
| JEB00022 | 5.90 | 0.00 | 33.94 | 39.00 | 29.96 | 34.03 |
| JEB00023 | 54.00 | 0.02 | 13.66 | 39.00 | 0.00 | 34.03 |
| JEB00024 | 12.21 | 0.02 | 46.10 | 39.00 | 17.24 | 34.03 |
| JEB00025 | 30.24 | 0.02 | 4.19 | 39.00 | 66.11 | 34.03 |
| JEB00028 | 77.30 | 0.02 | 16.30 | 0.00 | 38.21 | 34.03 |
| JEB00029 | 12.71 | 0.02 | 42.78 | 39.00 | 0.00 | 34.03 |
| JEB00030 | 0.97 | 0.02 | 17.43 | 39.00 | 0.00 | 34.03 |
| JEB00031 | 11.37 | 0.02 | 36.12 | 39.00 | 0.00 | 34.03 |
| JEB00032 | 83.98 | 0.17 | 13.88 | 39.00 | 0.00 | 34.03 |
| JEB00035 | 11.69 | 0.17 | 25.15 | 39.00 | 65.34 | 34.03 |
| JEB00036 | 32.79 | 0.17 | 20.43 | 39.00 | 44.11 | 34.03 |
| JEB00037 | 1.53 | 0.17 | 92.80 | 39.00 | 0.00 | 34.03 |
| JEB00040 | 54.97 | 0.17 | 19.03 | 0.00 | 19.09 | 34.03 |
| JEB00041 | 22.48 | 0.17 | 52.49 | 39.00 | 64.35 | 34.03 |
| JEB00042 | 81.15 | 0.17 | 12.20 | 39.00 | 49.57 | 34.03 |
| JEB00045 | 18.94 | 1.66 | 58.98 | 39.00 | 0.00 | 34.03 |
| JEB00046 | 56.17 | 1.66 | 11.58 | 39.00 | 10.34 | 34.03 |
| JEB00052 | 12.71 | 1.66 | 61.60 | 39.00 | 46.74 | 34.03 |
| JEB00065 | 53.66 | 1.66 | 21.90 | 39.00 | 0.00 | 34.03 |
| JEB00090 | 12.80 | 1.66 | 20.52 | 0.00 | 12.49 | 34.03 |
| JEB00113 | 0.40 | 1.66 | 30.21 | 39.00 | 32.90 | 34.03 |
| JEB00117 | 17.33 | 1.66 | 75.89 | 39.00 | 0.00 | 34.03 |
| JEB00122 | 59.77 | 16.61 | 29.59 | 39.00 | 13.04 | 34.03 |
| JEB00125 | 29.75 | 16.61 | 43.12 | 39.00 | 29.71 | 34.03 |
| JEB00126 | 80.07 | 16.61 | 82.83 | 39.00 | 15.78 | 34.03 |
| JEB00128 | 38.29 | 16.61 | 23.25 | 39.00 | 47.56 | 34.03 |
| JEB00133 | 24.44 | 16.61 | 18.76 | 39.00 | 231.90 | 34.03 |
| JEB00140 | 59.55 | 16.61 | 5.84 | 39.00 | 0.00 | 34.03 |
| JEB00144 | 7.02 | 16.61 | 43.00 | 39.00 | 0.00 | 34.03 |
| JEB00153 | 21.05 | 166.12 | 21.10 | 39.00 | 25.83 | 34.03 |
| JEB00169 | 21.26 | 166.12 | 43.07 | 0.00 | 35.83 | 34.03 |
| JEB00173 | 100.42 | 166.12 | 101.61 | 39.00 | 6.19 | 34.03 |
| JEB00174 | 44.96 | 166.12 | 36.63 | 39.00 | 56.75 | 34.03 |
| JEB00229 | 31.39 | 166.12 | 25.81 | 39.00 | 0.00 | 34.03 |
| JEB00254 | 79.39 | 166.12 | 114.89 | 39.00 | 45.92 | 34.03 |
| JEB00274 | 7.20 | 166.12 | 13.17 | 39.00 | 53.74 | 34.03 |

**Table S4.** Strain coverages in mock *E. lenta* distributions.

| StrainID | Exponential | Log step | Log-normal | Missing Strain | Zero inflated log-normal | Uniform |
| --- | --- | --- | --- | --- | --- | --- |
| JEB00079 | 80.48 | 0.00 | 23.89 | 94.28 | 139.19 | 77.14 |
| JEB00067 | 10.06 | 0.00 | 75.27 | 94.28 | 71.80 | 77.14 |
| JEB00066 | 14.50 | 0.04 | 88.01 | 0.00 | 10.12 | 77.14 |
| JEB00068 | 132.77 | 0.04 | 35.42 | 94.28 | 0.00 | 77.14 |
| JEB00069 | 30.02 | 0.04 | 119.53 | 94.28 | 104.47 | 77.14 |
| JEB00070 | 74.35 | 0.04 | 10.86 | 0.00 | 186.78 | 77.14 |
| JEB00080 | 190.05 | 0.38 | 42.27 | 94.28 | 86.06 | 77.14 |
| JEB00088 | 31.25 | 0.38 | 110.91 | 94.28 | 10.06 | 77.14 |
| JEB00082 | 2.38 | 0.38 | 45.20 | 94.28 | 0.00 | 77.14 |
| JEB00081 | 27.95 | 0.38 | 93.66 | 94.28 | 357.17 | 77.14 |
| JEB00077 | 206.48 | 3.82 | 35.98 | 94.28 | 74.48 | 77.14 |
| JEB00085 | 28.74 | 3.82 | 65.21 | 94.28 | 76.55 | 77.14 |
| JEB00083 | 80.62 | 3.82 | 52.97 | 0.00 | 76.00 | 77.14 |
| JEB00086 | 3.75 | 3.82 | 240.63 | 94.28 | 23.11 | 77.14 |
| JEB00072 | 135.15 | 38.18 | 49.35 | 94.28 | 79.58 | 77.14 |
| JEB00073 | 55.29 | 38.18 | 136.11 | 94.28 | 82.98 | 77.14 |
| JEB00074 | 199.54 | 38.18 | 31.63 | 0.00 | 17.91 | 77.14 |
| JEB00078 | 46.56 | 38.18 | 152.92 | 94.28 | 45.08 | 77.14 |
| JEB00084 | 138.10 | 381.83 | 30.02 | 94.28 | 0.00 | 77.14 |
| JEB00075 | 31.25 | 381.83 | 159.73 | 94.28 | 89.21 | 77.14 |
| JEB00076 | 131.93 | 381.83 | 56.79 | 94.28 | 100.41 | 77.14 |
| JEB00087 | 31.48 | 381.83 | 53.20 | 94.28 | 82.90 | 77.14 |

## Notes

https://github.com/BisanzLab/StrainR2

### Competing Interest Statement

The authors have declared no competing interest.

